# A method for quantifying *Phytophthora* oospore viability using fluorescent dyes and automated image analysis

**DOI:** 10.1101/2021.10.17.464154

**Authors:** Michael J. Fairhurst, Jochem N.A. Vink, Julie R. Deslippe, Monica L. Gerth

## Abstract

*Phytophthora* are eukaryotic microbes that cause disease in a wide range of agriculturally and ecologically important plants. During the *Phytophthora* disease cycle, thick-walled oospores can be produced via sexual reproduction. These resting spores can survive in the soil for several years in the absence of a host plant, thus providing a long-term inoculum for disease. The ability to quantitatively evaluate oospore viability is an important part of many phytopathology studies. Here, we tested six fluorescent viability dyes for their ability to differentially stain *Phytophthora agathidicida* oospores: SYTO 9, FUN-1, fluorescein diacetate (FDA), 5-carboxyfluorescein diacetate (CFDA), propidium iodide, and TOTO-3 iodide. Each dye was first tested individually with untreated or heat-treated oospores as proxies for viable and non-viable oospores, respectively. SYTO9, FUN-1, CFDA and propidium iodide stained untreated and heat-treated oospores indiscriminately. In contrast, FDA (a green-fluorescent viable cell stain) and TOTO-3 (a red-fluorescent non-viable cell stain) differentially stained untreated or heat-treated oospores with no cross-fluorescence. We then tested the efficacy of dual viability staining and in conjunction with a pipeline for automated image analysis. To validate the method, untreated and heat-treated oospores were mixed at specific ratios, dual-stained, and analyzed using the pipeline. Linear regression of the resulting data showed a clear correlation between the expected and measured oospore ratios (*dy/dx*=0.95, *R*^*2*^=0.88). Overall, the combination of dual-fluorescence staining and automated image analysis provides a high-throughput method for quantitatively assessing oospore viability and therefore can facilitate further studies on this key part of the *Phytophthora* disease cycle.

## Introduction

Viability staining is an essential technique in the study of plant pathogens: it is widely used to evaluate susceptibility to anti-microbial drugs, monitor disinfection protocols, and/or provide fundamental insights into pathogen physiology (Tian et al. 2019; Braissant et al. 2020). Methods for the viability staining of bacterial and fungal pathogens are well-established (Kwolek-Mirek et al. 2014; Kumar et al. 2019; Wurster et al. 2019). However, robust methods for assessing viability are lacking for many other microorganisms (Braissant et al. 2020).

*Phytophthora* are eukaryotic microorganisms that resemble filamentous fungi but are actually members of the class Oomycota and therefore are more closely related to diatoms and brown algae (Tyler et al. 2006). There are over 100 known species of *Phytophthora*, and many are notorious for their devastating impacts on crops and natural ecosystems (Kovacs et al. 2011; Kamoun et al. 2015; Drake et al. 2017; Corcobado et al. 2020; Susan et al. 2020). During the *Phytophthora* disease cycle, thick-walled oospores can be produced via sexual reproduction; some species of *Phytophthora* can also produce similar thick-walled chlamydospores via asexual reproduction (Erwin et al. 1996; Judelson et al. 2005). Once matured, these durable ‘survival spores’ can persist in soil and provide a long-term inoculum for disease (Judelson et al. 2005). Consequently, methods for the study of *Phytophthora* oospores are of great interest (Montes-Borrego et al. 2009; Bradshaw et al. 2020).

The goal of this study was to develop an effective viability staining protocol for *Phytophthora* oospores that is both quantitative and amenable to automated data analysis. The model organism for our study was *Phytophthora agathidicida*, a clade 5 *Phytophthora* species and a pathogen of kauri trees (*Agathis australis*) in New Zealand (Gadgil 1974; Beever et al. 2009; Waipara et al. 2013). *P. agathidicida* is homothallic and produces oospores readily via self-fertilization (Weir et al. 2015).

Existing methods for determining *Phytophthora* oospore viability primarily rely on methylthiazolyldiphenyl-tetrazolium bromide (MTT) or trypan blue staining (Cohen 1984; Beakes et al. 1986; Etxeberria et al. 2011). When stained with MTT, viable oospores may appear blue, rose/purple, or reddish-brown – while non-viable oospores may appear either black or unstained. Subtle color differences can be difficult to distinguish by human eye and/or automated image analysis (Meier et al. 1993; Etxeberria et al. 2011). Multiple studies have found MTT staining of oospores to be subjective, unstable, and have a high rate of false positives (Etxeberria et al. 2011; Williams 2015). Similar issues have been observed with trypan blue staining (Cohen 1984). In addition, trypan blue assays are time-sensitive; over time all cells will stain blue (regardless of viability), thus resulting in false negatives (Kwok et al. 2004; Kunjeti et al. 2016; Chan et al. 2020). Besides staining, another method sometimes used for determining viability is the direct plating of oospores on agar (Etxeberria et al. 2011).

However, due to the inherent dormancy of oospores, the germination rates can be very low *in vitro* (*e*.*g*. <10%) and conditions for germination vary for different species (Ribeiro et al. 1975; Flier et al. 2001; Xavier et al. 2010). Germination *in vitro* is further complicated for emerging pathogens such as *P. agathidicida* where the chemical stimulants and/or physical conditions required to break dormancy are not fully understood.

In terms of viability staining, fluorescent dyes offer key advantages (Kamiloglu et al. 2020). For example, fluorescent dyes provide more sensitivity and a wider dynamic range than colorimetric dyes (Kamiloglu et al. 2020). Multiple fluorescent dyes can also be used simultaneously (*e*.*g*. dual staining) which can improve accuracy (Tian et al. 2019). Lastly, fluorescent dyes are compatible with automated image analysis which can reduce user subjectivity and increase throughput (Chan et al. 2012).

In the present study, we tested six fluorescent dyes designed to quantify either the total number of oospores (*i*.*e*. SYTO 9 and FUN-1), the viable oospores (*i*.*e*. fluorescein diacetate (FDA) and 5-carboxyfluorescein diacetate (CFDA)), or the non-viable oospores (*i*.*e*. propidium iodide and TOTO-3 iodide). Each dye was assessed individually, and then the best two dyes were combined for dual viability staining. We assessed their suitability for this method by investigating to what extent these dyes can discriminate between untreated and heat-treated (as proxies for viable and non-viable) oospores and whether they fluoresce in a single color or both red and green fluorescence channels, which would hamper the use of dual color staining.

SYTO 9 is a cell-permeant, nucleic acid dye that is commonly used for assessing total cell counts of bacteria and yeast (Jin et al. 2005; McGoverin et al. 2020). SYTO 9 enters both viable and non-viable cells and binds to DNA/RNA, resulting in green fluorescence. FUN-1 is also designed to stain all cells within a population, but it is a multi-color fluorescent probe (Millard et al. 1997). It passively diffuses into cells, initially producing green fluorescence (Millard et al. 1997). Subsequent processing by metabolically active cells results in red fluorescence, and dead cells fluoresce yellow-green (Millard et al. 1997). FUN-1 was originally developed as a vital stain for yeast (Millard et al. 1997), but more recently has also been used to assess *Phytophthora capsici* sporangia and zoospores (Lewis Ivey et al. 2014).

The viable cell stains FDA and its derivative CFDA are both non-fluorescent molecules that can diffuse freely through cell membranes (Rotman et al. 1966). In live cells, FDA is hydrolyzed by intracellular esterases to liberate the green-fluorescent product fluorescein (Rotman et al. 1966). Fluorescein accumulates in viable cells with intact membranes, and therefore the resulting green fluorescence is a marker of both active metabolism and an intact cell membrane (Rotman et al. 1966). CFDA operates by the same mechanism but is more negatively charged compared to FDA and is therefore typically better retained in cells (Goodall et al. 1982; Breeuwer et al. 1995; Braissant et al. 2020). FDA/CFDA are often used in conjunction with red-fluorescent dyes such as propidium iodide or TOTO-3 for dual viability staining (Braissant et al. 2020).

PI and TOTO-3 are membrane-impermeant to viable cells. However, in non-viable cells with disrupted membranes, propidium iodide and TOTO-3 can enter, bind to DNA and RNA, and produce red fluorescence (Braissant et al. 2020).

Here, we show that FDA and TOTO-3 can be used to quantify viable and non-viable oospores, respectively. We have also developed a pipeline for automated analysis of the resulting fluorescence microscopy images. Used together, these methods facilitate the rapid and accurate assessment of oospore viability.

## Materials and Methods

### Materials

*P. agathidicida* isolates NZFS 3770 and NZFS 3772 were provided by Scion (New Zealand Forest Research Institute, Rotorua, New Zealand). Genome sequences for both isolates are available (Studholme et al. 2016; Cox et al. 2022). SYTO 9, FUN-1, CFDA, TOTO-3, and PI were sourced from Thermo Fisher Scientific (Waltham, Massachusetts). FDA was sourced from Sigma-Aldrich (St. Louis, Missouri). Fluorescent dyes were supplied or prepared as dimethylsulfoxide (DMSO) stocks and stored at -20 °C in small aliquots protected from light. Stock solutions were diluted with sterile water to achieve the final working concentrations immediately before use. The final percentage of DMSO was <2% in all assays.

### Oospore production

*P. agathidicida* oospores were prepared as described previously (Lacey et al. 2021). Briefly, cultures were grown in 4% w/v clarified V8 broth supplemented with 30 *µ*g/ml β-sitosterol (Sigma-Aldrich, St. Louis, Missouri) and incubated in the dark at 22 °C for 4 to 6 weeks. The resulting mycelial mats were homogenized, sonicated, and then filtered using 100 *µ*m and 40 *µ*m Greiner Bio-One EASYstrainers (Frickenhausen, Germany). The oospore suspension was then pelleted at 1,200 × g for 10 min, the supernatant discarded, and the oospores resuspended in 5 ml of sterile water. Oospore numbers were estimated using disposable hemocytometers. In initial single-dye experiments, heat-treated oospores were produced by incubating the purified oospores 98 °C for 1h. In subsequent dual-stain and time course experiments, heat treatment was applied for up to 24 h, because we observed a slightly higher percentage of non-viable oospores with longer heat treatment.

### Microscopy

Oospores were imaged using an Olympus BX63 fluorescence microscope with a 4× or 10× objective (and a 0.63× reductive camera lens) using brightfield and/or fluorescence microscopy. Green fluorescence was measured using Chroma filter 31001 (480 nm excitation center, 30 nm bandwidth; 535 nm emission center, 40 nm bandwidth) and an. Red fluorescence. was measured using Chroma filter 49008 (560 nm excitation center, 40 nm bandwidth; 630 nm emission center, 75 nm bandwidth) with a 50 ms exposure.

### Oospore germination

The germination rates of untreated and heat-treated (98 °C for 24 h) oospores were assessed by plating oospores on water agar plates amended with 1% w/v kauri extract to stimulate germination. After 4 d of incubation at 22 °C, the percentages of germinated oospores were assessed visually at 40x magnification. An oospore was considered germinated if a germ tube at least twice the length of the oospore diameter was present. Three independent replicates were performed with one hundred oospores visually assessed per replicate.

### Oospore viability staining

Each dye was first tested individually on untreated and heat-treated (at 98 °C for 1 h) *P. agathidicida* NZFS 3770 oospores. Oospores (∼2,500) were pelleted (1,200*g* for 10 min) and the supernatant was removed. Oospores were then resuspended in 10 μl of each dye at its final working concentration. Each dye was screened to optimize the concentrations, incubation temperatures, and incubation times. SYTO 9, FDA, and PI were tested at 2 *µ*M, 20 *µ*M, and 200 *µ*M; FUN-1 was tested at 8 *µ*M, 40 *µ*M, and 200 *µ*M; and TOTO-3 was tested at 1 *µ*M, 10 *µ*M, and 100 *µ*M). Each concentration of dye was assessed at both 22 and 37 °C and for three different incubation times (4 h, 24 h, and 48 h). The oospores were imaged, and the best conditions were chosen qualitatively based on the ability of each dye to differentially stain untreated/heat-treated oospores (Supplementary Figures S1-S3). Three independent replicates were performed for each condition tested. Based on these results, the selected conditions for subsequent experiments were as follows: 2 *µ*M SYTO 9 at 37 °C for 4 h; 20 *µ*M PI at 37 °C for 4 h; 10 *µ*M TOTO-3 at 37 °C for 4 h; 40 *µ*M FUN-1 37 °C for 24 h; 200 *µ*M FDA at 37 °C for 24 h and 200 *µ*M CFDA at 37 °C for 24 h).

Dual staining of oospores was conducted using FDA and TOTO-3. Oospores from two different *P. agathidicida* isolates (NZFS 3770 and NZFS 3772) were tested. Heat-treated oospores were prepared at 98 °C for 0–24 h. For dual staining, the oospores were first stained with 200 μM FDA for 20 h at 37 °C. Then, TOTO-3 was added to a concentration of 20 μM and the samples were incubated for a further 4 h at 37 °C before imaging. Three independent replicates were performed.

### Image analyses

For high-throughput image analyses, we combined a set of image and data analysis algorithms. Firstly, we used the interactive machine learning software Ilastik (Berg et al. 2019) to automatically identify oospores in brightfield images. For optimal oospore segregation, two different pixel classifiers were trained, one to identify oospore pixels in general and one for edge detection. Second, a pipeline was written in CellProfiler (McQuin et al. 2018) which identified oospore objects from the two-pixel classification images using a watershed algorithm. Within the same pipeline, oospore red- and green-fluorescence intensities were extracted and oospores were classified as fluorescent if their upper quartile intensity was above a set threshold (0.1 for the green channel and 0.08 for the red channel). The upper-quartile intensity is the intensity value of the pixel for which 75% of the pixels in the object have lower values and was used as it is less sensitive to variability caused by over- or under-estimation of the oospore region. Data on the size and fluorescence intensity for each oospore were stored as outputs. Lastly, these data were further processed and visualized in R v3.6.0, with Rstudio v1.2.5001 (R Core Team 2017) in combination with the ggplot2 package for graphic generation (Wickham 2016). The complete CellProfiler pipeline and R scripts used for analyses are provided at https://github.com/MMELab/Oospore-viability.

### Statistics

To investigate the significance of differences observed between treated and untreated samples stained by individual dyes, we applied an arc-sine square root transformation to the fraction of either green or red-stained oospores and subsequently ran the Student’s t-test. To investigate the significance of differences observed between *P. agathidicida* isolates 3770 and 3772 in our time course experiment, we used a mixed analysis of variance (ANOVA) model with isolates as the between variable and time as the within variable. The dependent variable was the fraction of green oospores transformed with the arc-sine square root transformation. All statistics were performed in R and we used α=0.05 for all statistical tests.

## Results

### Viability staining with individual dyes

Each dye was assessed individually for its ability to distinguish between untreated and heat-treated oospores and also for its potential compatibility with complementary dyes (*e*.*g*. no cross-fluorescence).

SYTO-9 (a green-fluorescent nuclei acid dye that reports the total population of cells, regardless of viability) stained both untreated and heat-treated oospores resulting in green fluorescence (Fig. 1A, 1D). However, particularly for untreated oospores, a significantly (*t* = 7.7, *p* = 0.01) larger fraction (87%) remained unstained (Fig 1C). No red cross-fluorescence was observed with SYTO 9 (Fig. 1B, 1E).

**Figure 1.**
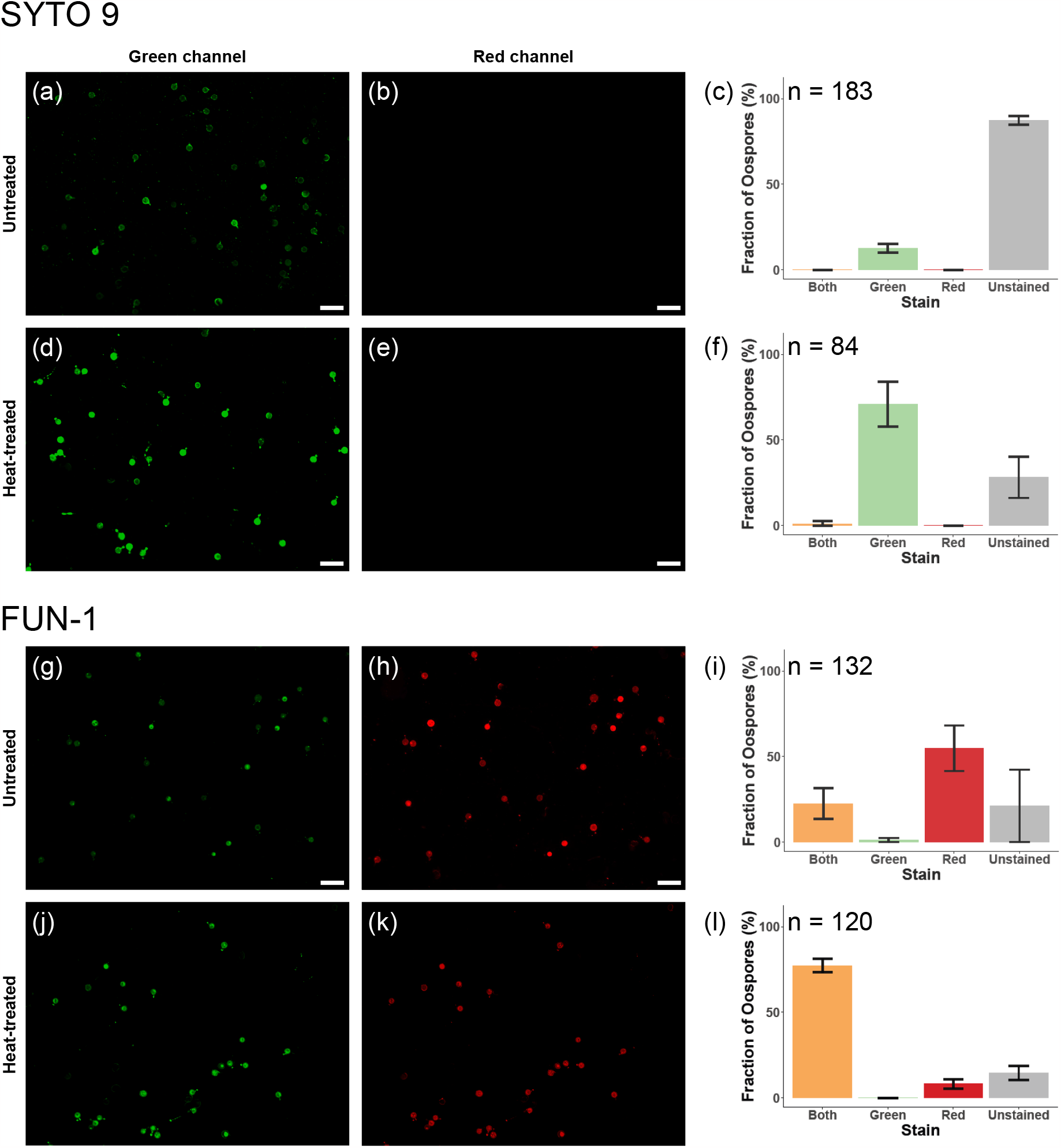
Comparison of two universal (total cell count) dyes: SYTO 9 and FUN-1. Untreated and heat-treated (1 h at 98 °C) P. agathidicida NZFS 3770 oospores were stained with SYTO 9 **(a-f)** or FUN-1 **(g-l)**, then imaged with both green and red filters, and the fractions of stained oospores classified using the automated pipeline. Each image is representative of three independent experiments. Scale bar = 100 μm. Plots represent oospore counts from three independent experiments. The total number of oospores counted (n) per plot are as follows: n=183 **(c)**; n=84 **(f)**; n=132 **(i)**; n=120 **(l)**. Error bars represent standard deviations.

The other total population dye tested, FUN-1, caused a fraction of oospores to fluoresce both green and red, regardless of treatment (Fig. 1G–1L). For heat-treated oospores, there was no significant difference in the proportion of red oospores (from 85 to 77%; *t* = 0.096, *p* = 0.93), however, there was a significant increase in green oospores (from 24% to 77%; *t* = 9.3, *p* = 0.0031) upon heat treatment.

The viable cell stain FDA produced green fluorescence with untreated oospores (Fig. 2A-C). No green fluorescence was observed with heat-treated oospores (Fig. 2D-F). The difference between untreated and heat-treated oospores is significant (*t* = -13, *p* = 0.002), and no red cross fluorescence was observed for either sample (Fig. 2B-C, E-F). A proportion (47%) of the untreated oospores were classified as unstained.

**Figure 2.**
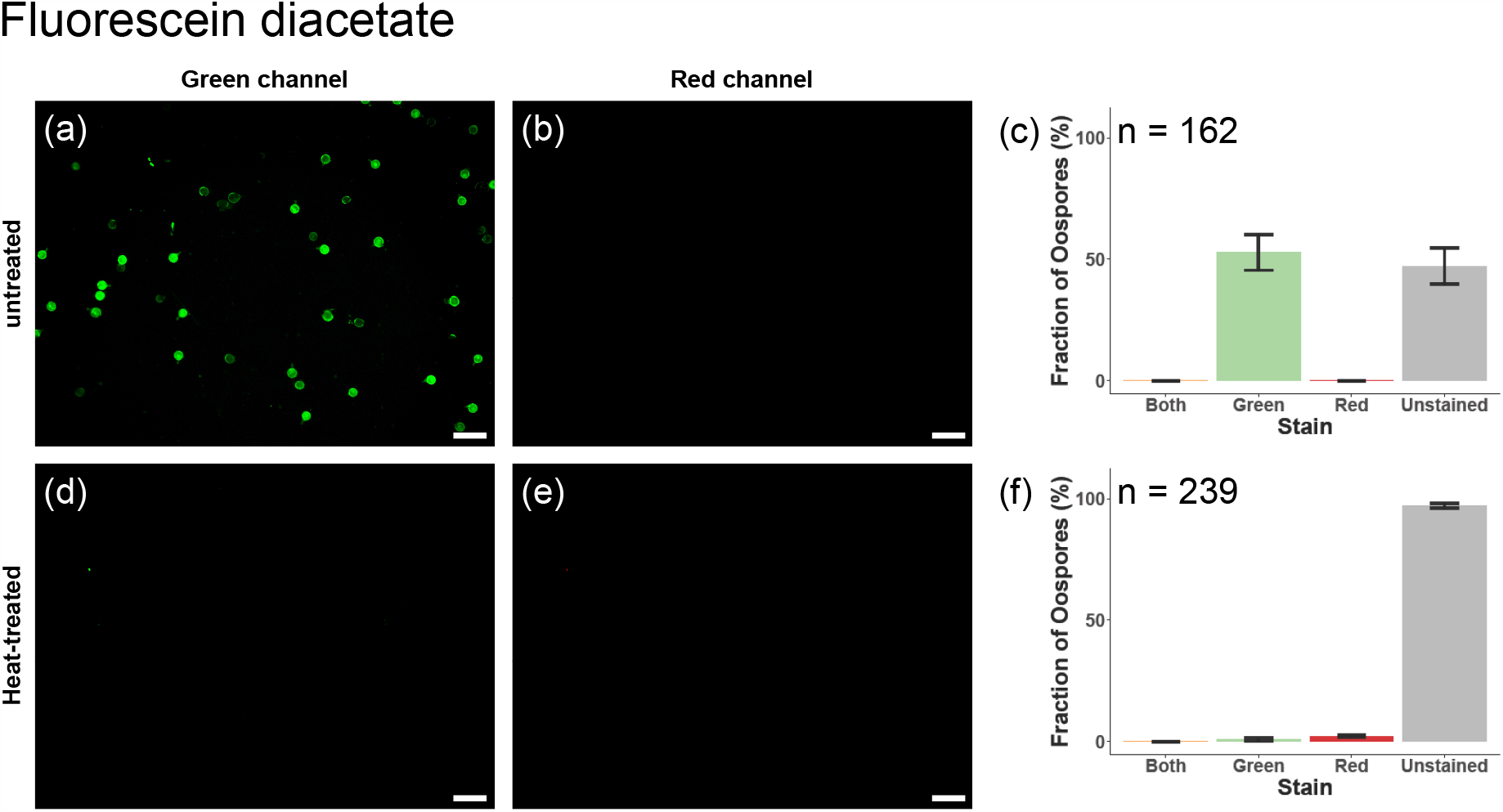
Assessment of fluorescein diacetate (FDA) as a potential dye for the detection of viable oospores. Untreated and heat-treated (1h at 98 °C) P. agathidicida NZFS 3770 oospores were stained with FDA, then imaged with both green and red filters, and the fractions of stained oospores were classified using the automated pipeline. **(a-c)** untreated oospores and **(d-f)** heat-treated oospores. Each image is representative of three independent experiments. Scale bar = 100 μm. Plots represent total oospore counts from three independent experiments. The total number of oospores counted (n) per plot are as follows: n=162 **(c)**; n=239 **(f)**. Error bars represent standard deviations.

We also tested whether the FDA derivative – CFDA – would further improve the signal to background detection of untreated versus heat-treated oospores. However, we found in a direct comparison of their intensities that the FDA dye provided a better distinction between untreated and heat-treated oospores (Supplementary Figure S4). We, therefore, determined FDA to be the better candidate to take forward for further testing.

The two red fluorescent, non-viable cell stains tested were propidium iodide and TOTO-3 (Fig. 3). Propidium iodide was not able to distinguish between viable and non-viable oospores, with both types exhibiting red fluorescence (*t* = 1.2, *p* = 0.32; Fig 3A-F). In contrast, a majority (∼70%) of heat-treated oospores stained with TOTO-3 produced a red fluorescence signal (Fig. 3K-L) with little to no green cross-fluorescence (Fig. 3J). A fraction of the untreated oospores (∼33%, Fig. 3I) also exhibited red fluorescence (Fig. 3H,3I), but the difference between the treatments was significant (*t* = 6.6, *p* = 0.014).

**Figure 3.**
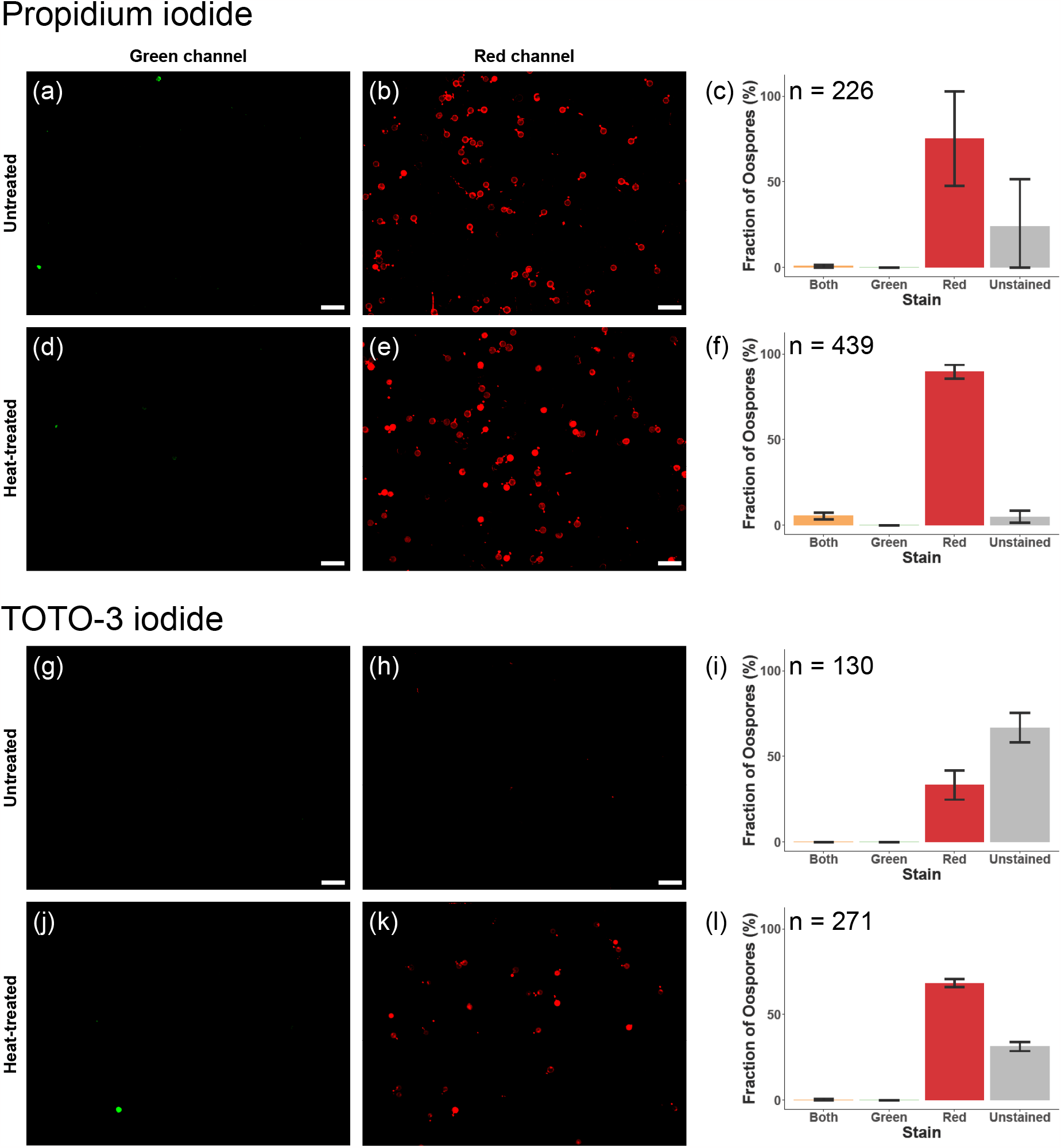
Comparison of dyes for the detection of non-viable oospores: Propidium iodide and TOTO-3. Untreated and heat-treated (1 h at 98 °C) P. agathidicida NZFS 3770 oospores were stained with propidium iodide **(a-f)** or TOTO-3 **(g-l)**, then imaged with both green and red filters, and the fractions of stained oospores classified using the automated pipeline. Each image is representative of three independent experiments. Scale bar = 100 μm. Plots represent the oospore counts from three independent experiments. The total number of oospores counted (n) per plot are as follows: n=226 **(c)**; n=439 **(f)**; n=130 **(i)**; n=271 **(l)**. Error bars represent standard deviations.

Overall, two dyes were identified that differentially stained oospores based on viability when used individually: FDA (a green-fluorescent viable cell stain) and TOTO-3 (a red-fluorescent non-viable cell stain).

### Oospore dual viability staining

Next, we tested the efficacy of the FDA and TOTO-3 when used together for dual viability staining (Fig. 4). With dual stained (NZFS 3770) oospores, ∼84% of the untreated oospores fluoresced green (Fig. 4A). This is more than the ∼6% that germinated after 4 d, however, germination potentially occurs over a much larger timeframe and may require unknown factors. A small fraction of oospores (1.5%) within the untreated oospore population also exhibited red fluorescence (Fig. 4A). In the dual-stained heat-treated (1hr at 98°C) oospore samples most cells produced a clear red fluorescence signal (82%), though several oospores (1.3%) appeared yellow (Fig. 4B). Both green and red fluorescent oospore fractions differed significantly between the two treatments (*t* = 16, *p* = 0.00041 and *t* = -11.411, *p* = 0.00015; green and red respectively).

**Figure 4.**
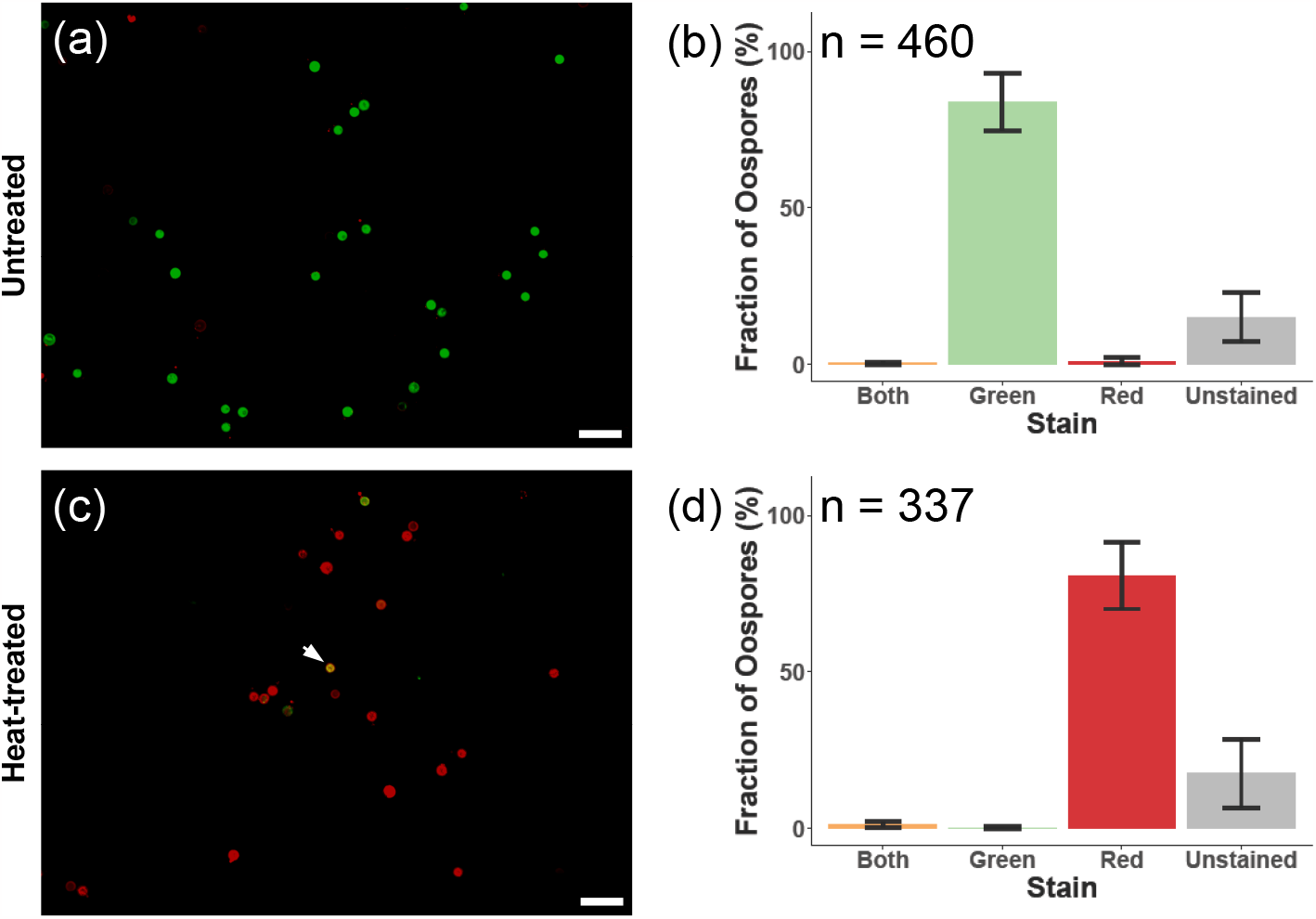
Dual viability staining of P. agathidicida NZFS 3770 oospores with FDA and TOTO-3. The merged images (green/red fluorescence) are shown for **(a & b)** untreated oospores and **(c & d)** heat-treated (1h at oospores 98 °C) and the fractions of stained oospores were classified using the automated pipeline. The arrow in panel (**c**) highlights a yellow oospore stained with both dyes. Images are representative of three independent experiments. Scale bar=100 μm. Error bars represent standard deviations.

We hypothesized that the yellow oospores represent a third state: damaged, but still metabolically active. To further explore this observation, we conducted a time course of oospore heat-inactivation (Fig. 5). The proportion of green (viable) and yellow (damaged) cells decreases steadily over the time course (Fig. 5). After 24 h of heat treatment, only non-viable (red) oospores are observed (Fig. 5). Similar results were obtained with a second isolate (NZFS 3772); however, the oospores of this isolate were significantly more robust (*F*(6,36) = 2.52, *p* = 0.038), with higher percentages of viable and damaged oospores surviving the heat treatment (Fig. 5).

**Figure 5.**
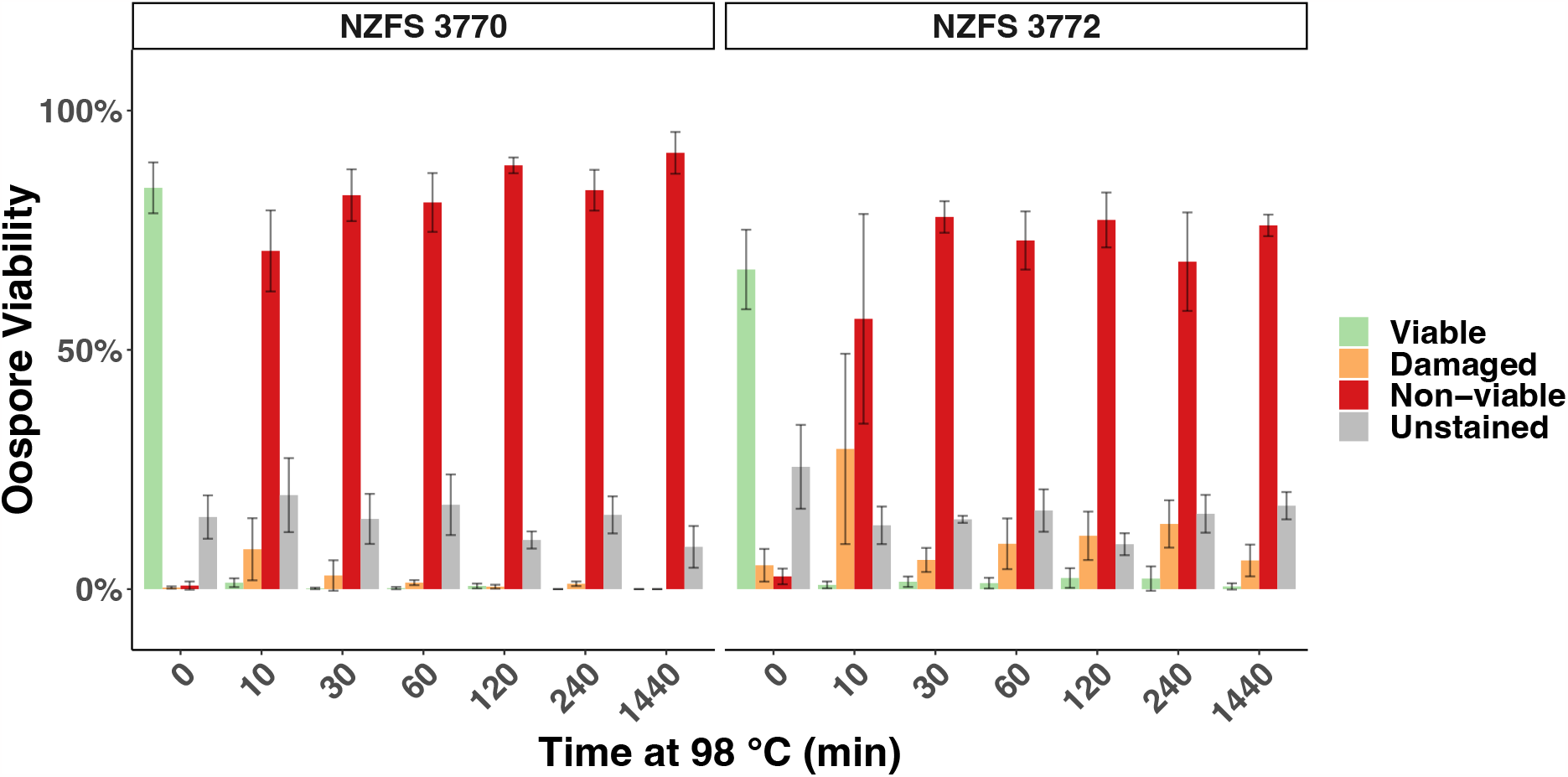
Dual-viability staining and time-course of oospore heat-inactivation. P. agathidicida NZFS 3770 and NZFS 3772 oospores were incubated at 98 °C for up to 24 h (1440 min). Samples were collected at the indicated time points, dual-stained with FDA and TOTO-3, and then imaged. The resulting data were analyzed in CellProfiler and plots were generated using R. FDA-stained (viable) oospores are shown as green bars; TOTO-3 stained (non-viable) oospores are shown as red bars; oospores stained by both dyes (damaged) are shown as orange bars. The bars represent the means of data from three independent experiments. Error bars represent the standard errors of the means. Significantly higher numbers of viable and/or damaged oospores of NZFS 3772 survived heat-treatment, as compared to NZFS 3770 (ANOVA F(6,36) = 2.52, p = 0.038)

### Automated pipeline for quantification of oospore viability

Finally, we sought to establish an automated image analysis pipeline that would improve the throughput of oospore viability assays. Our image analysis pipeline involved three steps: (i) total oospore counts were generated via machine learning analyses of the brightfield images (Supplementary Figure S5) (ii) CellProfiler was used to generate raw counts of viable (green), non-viable (red), and damaged (yellow) oospores (iii) an R-script was used to automatically transform the data, convert the total counts to percentages, and plot the data as a bar graph. To validate this pipeline, we dual-stained known ratios of untreated and heat-treated oospores (as proxies for viable and non-viable oospores) and analyzed the images. Overall, the experimentally determined proportions of untreated/heat-treated oospores matched well with the actual proportions (Fig. 6). The viability data could be fit using linear regression analysis (Fig. 6) and this regression showed a clear correlation between the expected and measured oospore ratios (*dy/dx*=0.95, *R*^*2*^=0.88).

**Figure 6.**
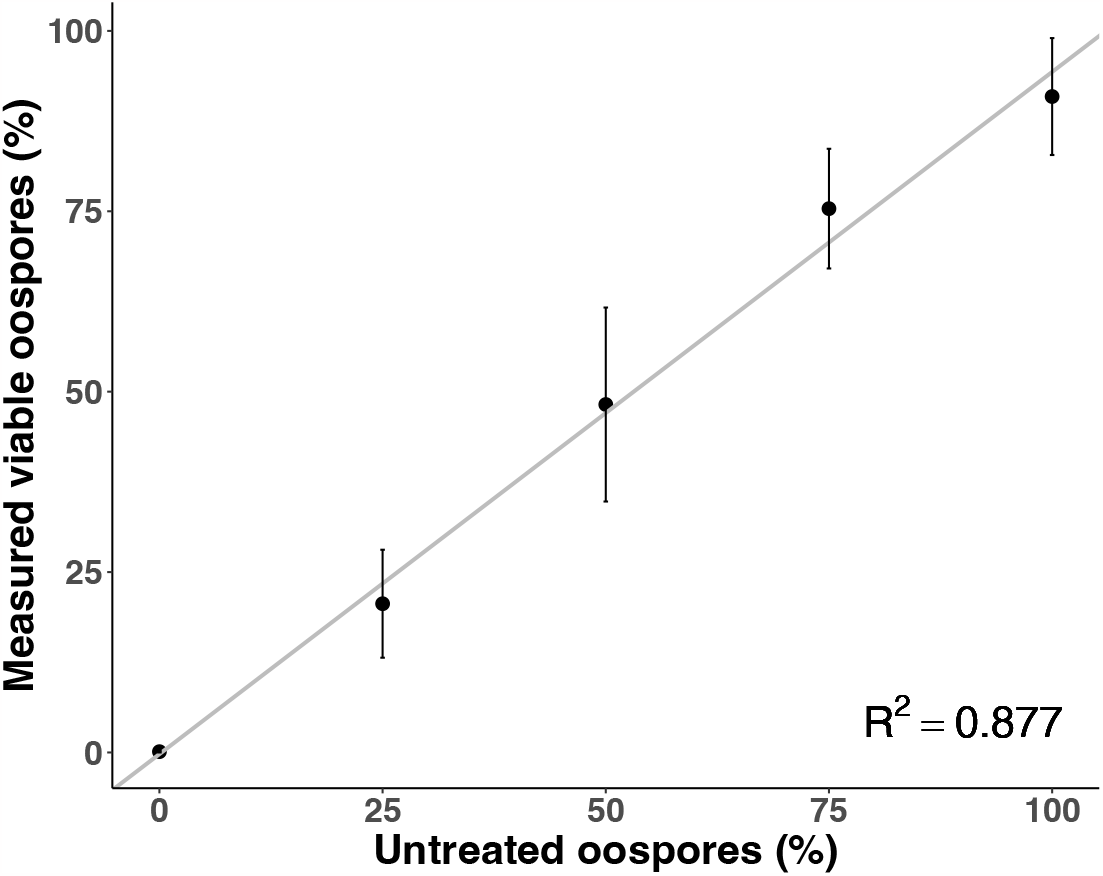
Measured viability of untreated and heat-treated oospores mixed at known proportions. Untreated and heat-treated (98 °C for 24 h) oospores were mixed in known proportions, dual stained with FDA and TOTO-3, and the stained oospores were counted and classified using the automated pipeline. The grey line represents the linear regression of the actual versus measured percentages of viable oospores. The data points represent the mean of three independent experiments; error bars represent the standard errors of the means.

## Discussion

In this study, we have established a fluorescence-based method for detecting the viability of *Phytophthora* oospores that can be combined with machine learning software for automated analysis. We identified two dyes – FDA and TOTO-3 – that can be used together for dual-viability staining. The use of multiple fluorescent dyes provides additional information as compared to single dyes (Tian et al. 2019). For example, in our single-dye assay using just fluorescein diacetate, a large proportion of oospores (47%) were classified as ‘unstained’ (Fig. 1). Upon manual inspection of brightfield images, we found that this unstained fraction contained a mixture of intact oospores (potentially non-viable) as well as partly broken oospores and empty shells (Supplementary Fig. 6). The addition of a second dye for non-viable cells (*e*.*g*. TOTO-3 iodide) allows this population to be further classified as ‘non-viable’ or unstained (*i*.*e*. debris). Dual staining, therefore, provides richer data, allowing oospores to be quantitatively classified as viable, non-viable, or unstained debris. Another advantage of using dual staining is that it also can potentially identify viable but damaged oospores (which appear yellow when dual-stained with FDA and TOTO-3). The ability to detect damaged oospores is not completely unexpected; similar intermediate states have been observed with live/dead staining of bacteria (Berney et al. 2007). Here, our results highlighted potential differences amongst isolates in the resistance of oospores to heat-treatment, with isolate NZFS 3772 having higher percentages of viable and damaged oospores surviving heat treatment (Fig. 5). Previous studies have established that there is variability in virulence among *P. agathidicida* isolates (Drake et al. 2017). However, potential differences in oospore robustness or resistance to treatments have not yet been explored; this will be an interesting avenue for future studies.

Ultimately, a key advantage of this new fluorometric method is its compatibility with automated image analysis. Our pipeline for automated image analysis increases both the throughput and reliability of viability assays – particularly in comparison to colorimetric dyes such as MTT where the color change is assessed subjectively (Etxeberria et al. 2011; Dick et al. 2013). These fluorescent dyes are also compatible with flow cytometry (Davey et al. 2020). Future studies could explore the use of this dual viability staining method with flow cytometry to further improve the reliability and throughput, allowing much larger populations to be assessed and/or sorted (Kummrow et al. 2013).

*Phytophthora* species remain some of the world’s most destructive plant pathogens. Our improved method for viability staining of oospores will facilitate further studies on this key part of the *Phytophthora* disease cycle. For example, this assay could be used to rapidly evaluate potential oospore biocidal treatments or to provide insights into the optimal conditions for oospore production, batch-to-batch variation, and/or conditions for survival. Ultimately, a better understanding of oospore viability can be used to develop tools and inform strategies for disease management.

## Acknowledgements

We thank Dr. Eiren Sweetman (School of Biological Sciences, Victoria University of Wellington) for assistance with oospore production and the CFDA staining experiments. We thank Dr. Lisa Woods (School of Mathematics and Statistics, Victoria University of Wellington) for assistance with the statistical analysis.

## Author Contributions

MJF: conceived and designed experiments; collected data; analyzed data; contributed to writing and editing of the manuscript. JNAV: collected data; analyzed data; contributed to writing and editing of the manuscript. JRD: supervision (secondary); contributed to review and editing of the manuscript. MLG: supervision (primary); funding acquisition; project administration; conceived and designed experiments; analyzed data; contributed to writing and editing of the manuscript.

## Code Availability

Instructions and the latest code for data processing and analysis using our pipeline can be found on the GitHub page https://github.com/MMElab/Oospore-viability.

