## Supplementary Materials for "A method for quantifying *Phytophthora* oospore viability using fluorescent dyes and automated image analysis"

bioRχiv

**Supplementary Figure S1**

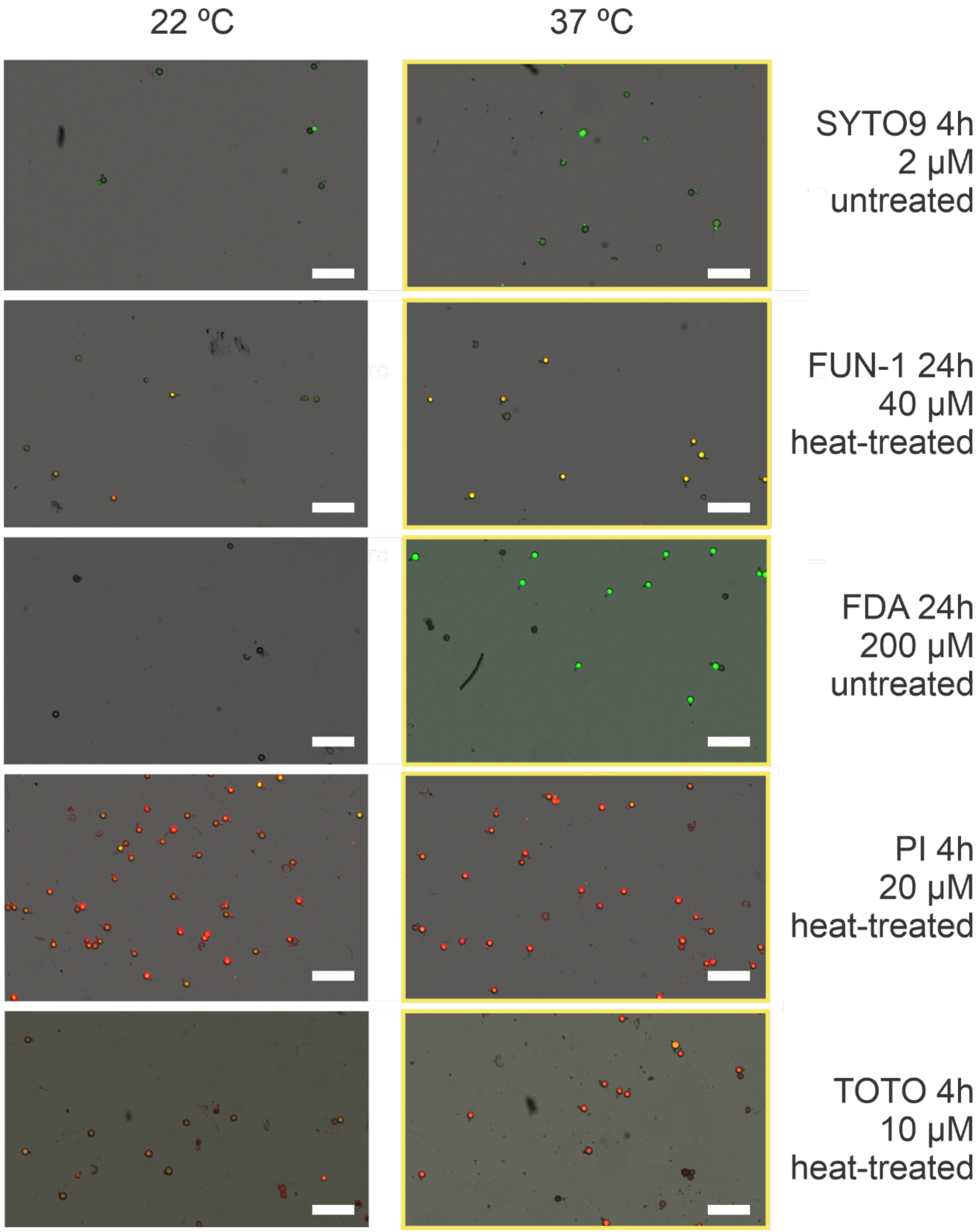


**Supplementary Figure S1. Optimisation of incubation temperature on untreated and heat-treated oospores.** Untreated and heat-treated oospores were tested with five dyes, at three concentrations (2, 20 and 200 $\mu$M for SYTO 9, FDA and PI; 8, 40 and 200 $\mu$M for FUN-1; and 1, 10 and 100 $\mu$M for TOTO-3 iodide), two temperatures (22 and 37 °C), and three incubation times (4, 24, and 48 h). These were imaged in the green (FITC) and red (Texas Red) channels and qualitatively evaluated to find the best condition (yellow outline) where the dyes stained oospores with the desired effect (either a total count or differentially stained for viability). Selection of data shown. Scale bar indicates 200 $\mu$m.

**Supplementary Figure S2.**

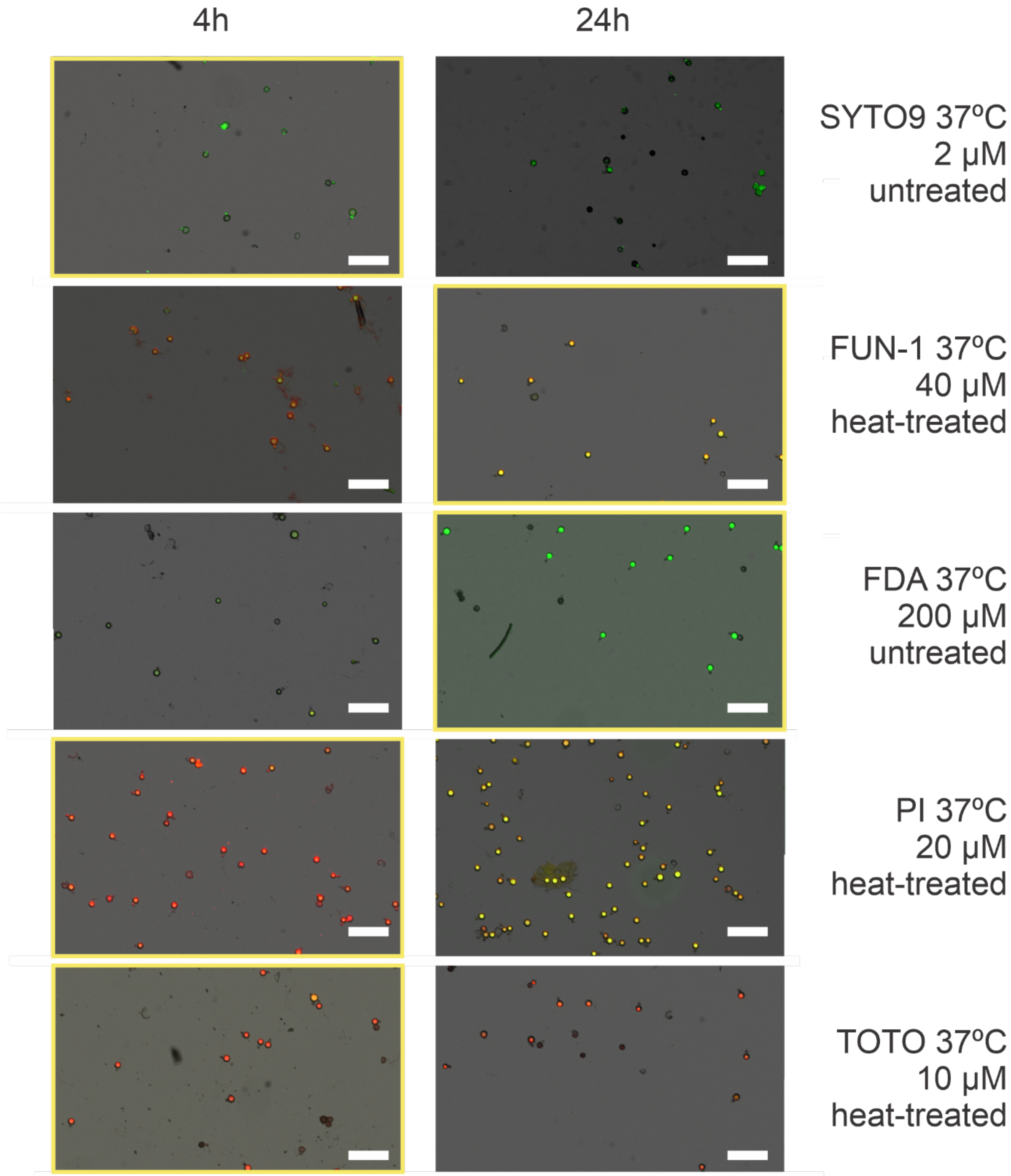


**Supplementary Figure S2. Optimisation of incubation time for five dyes on untreated and heat-treated oospores.** Other subset of the same experiment as Supplementary Figure S2. These were imaged in the green (FITC) and red (Texas Red) channels and qualitatively evaluated to find the best condition (yellow outline) where the dyes stained oospores with the desired effect (either a total count or differentially stained for viability). Scale bar indicates 200 $\mu$m.

**Supplementary Figure S3.**

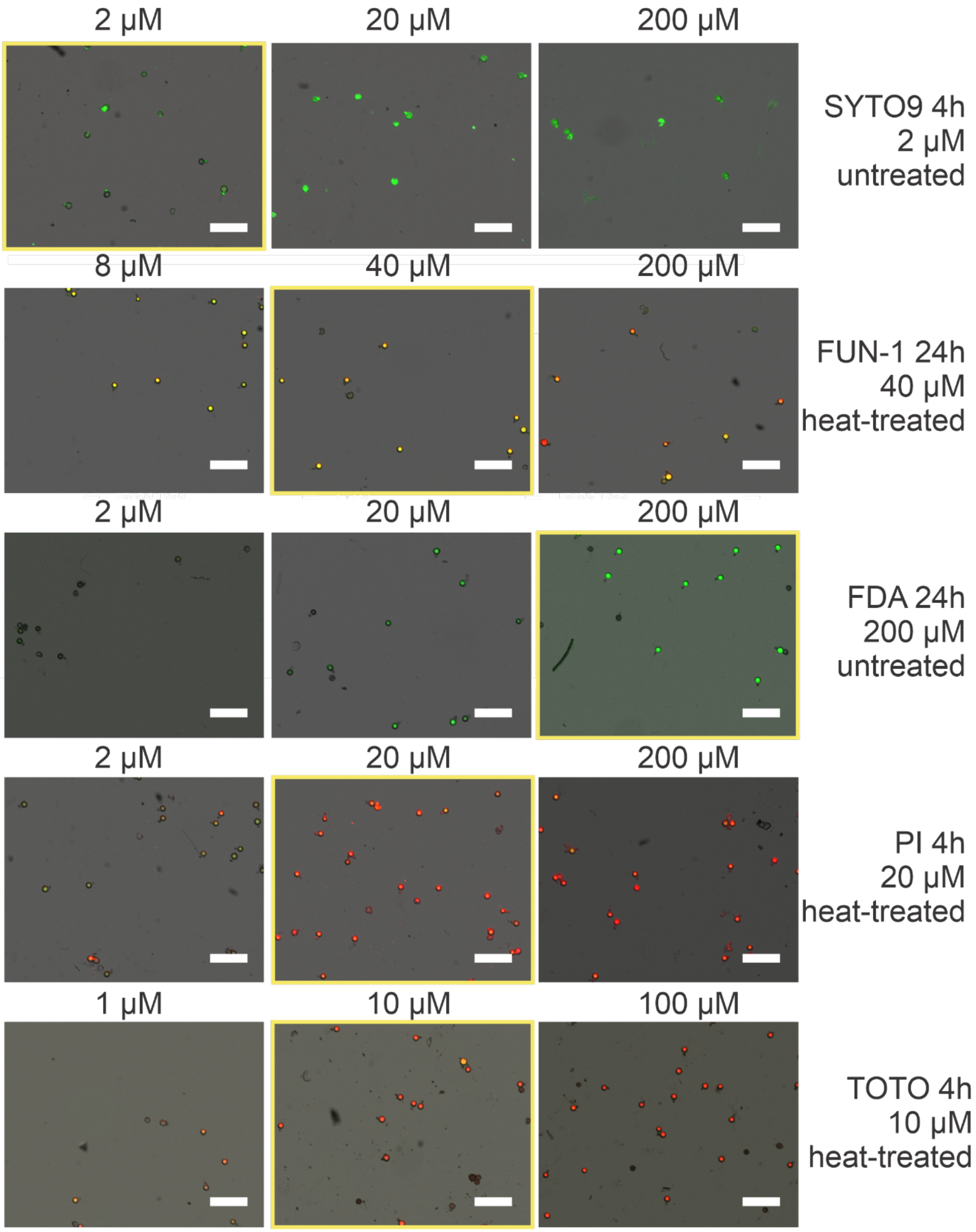


**Supplementary Figure S3. Optimisation of dye concentration on untreated and heat-treated oospores.** Other subset of the same experiment as Supplementary Figure S2. These were imaged in the green (FITC) and red (Texas Red) channels and qualitatively evaluated to find the best condition (yellow outline) where the dyes stained oospores with the desired effect (either a total count or differentially stained for viability). Scale bar indicates 200 $\mu$m.

**Supplementary Figure S4.**

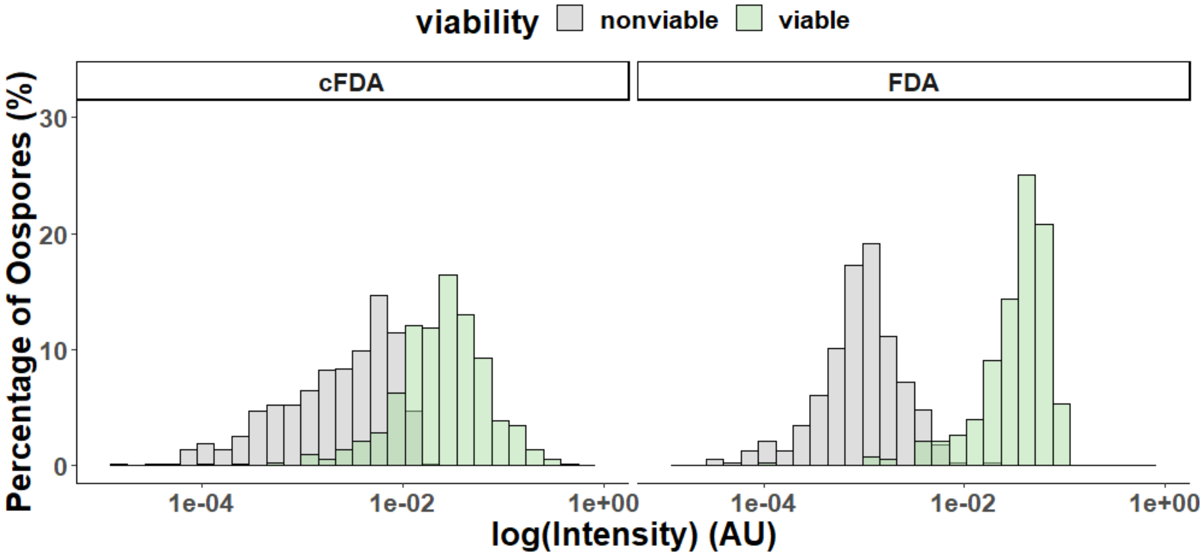


**Supplementary Figure S4. Intensity distributions of oospores stained with cFDA versus FDA.** A mixture of viable (green) and non-viable (grey) oospores were incubated with either cFDA (left) or FDA (right). The intensity of each oospore was determined as the 75% upper quartile of all pixels within the oospore region. The log(Intensity) values were binned into frequency histograms.

**Supplementary Figure S5.**

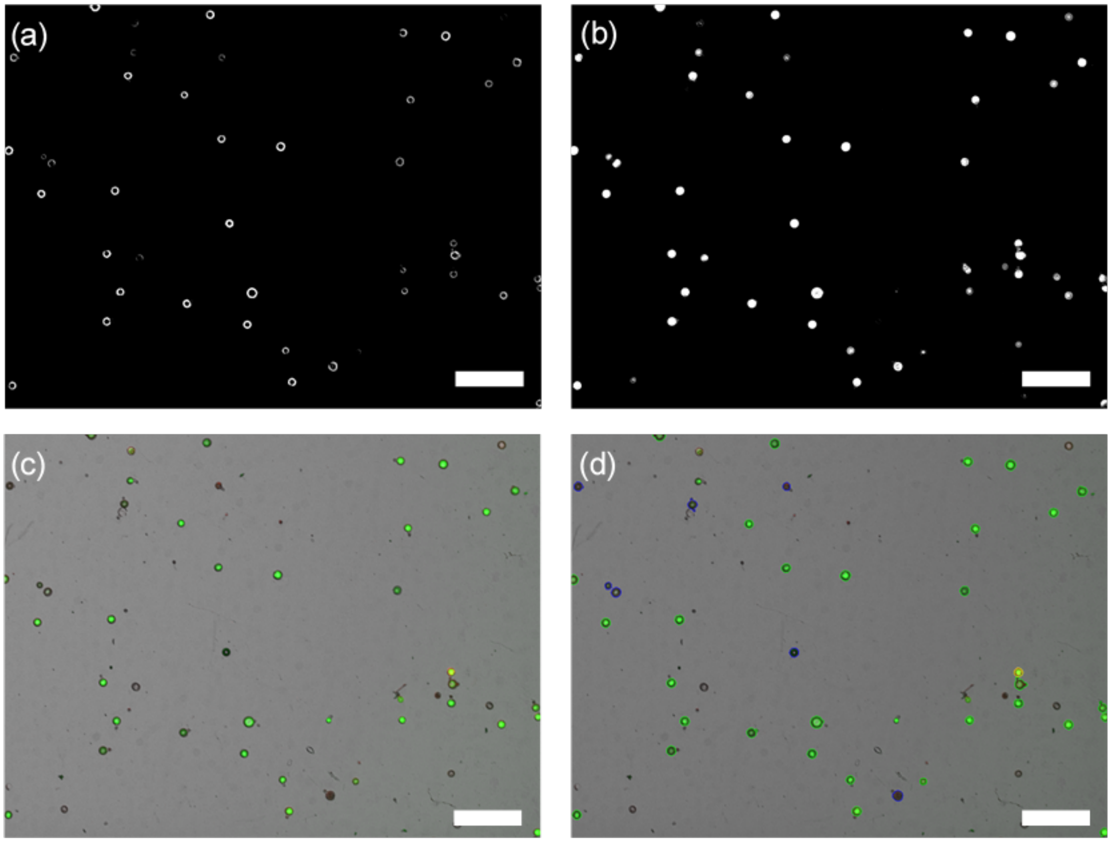


**Supplementary Figure S5. Stages in automated classification of dual-stained oospores using machine learning.** First, oospore regions are detected within a brightfield image from a combination of two machine learning algorithms, focused on either the edges (**a**) or oospores as a whole **(b)**. A combined brightfield and fluorescence image is then generated **(c).** This allows for the classification of oospores from the combined image **(d)** into either unstained (blue), single stained (green, red) or double stained (yellow) oospores. Scale bar = 200 µm.

**Supplementary Figure S6.**

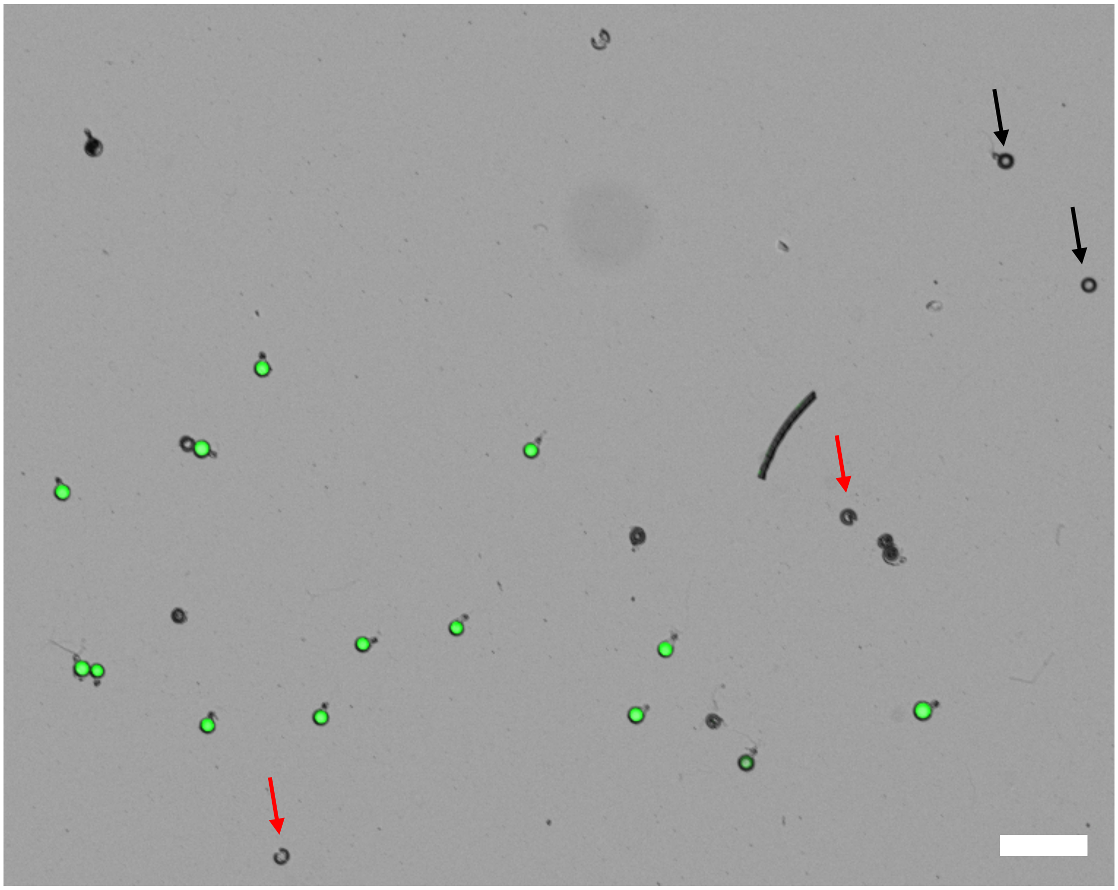

**Supplementary Figure S6. Morphology of unstained oospores.** Overlay of brightfield and fluorescence images of oospores stained with FDA (200 $\mu$M). The remaining fraction of unstained oospores differ in morphology, some have a clear compromised cell wall (red arrows), whereas others are indistinguishable from stained/healthy oospores (black arrows). Scale bar = 100 µm.
